# A tale of two lakes: divergent evolutionary trajectories of two *Daphnia* populations experiencing distinct environments

**DOI:** 10.1101/2024.04.19.590332

**Authors:** Matthew J. Wersebe, Torsten Günther, Philip K. Morton, Lawrence J. Weider, Dagmar Frisch

## Abstract

Most studies of local adaptation substitute the correlation between spatial distance and environmental heterogeneity for the temporal dynamics over which local adaptation evolves. The availability of detailed ecological and genomic information from lake sediments provides an opportunity to study local adaptation with unparalleled clarity from the temporal perspective. Inference can be further enhanced by including multiple lakes along ecological axes to further isolate the effects of ecological change in driving local adaptation. Lakes throughout the world face the impact of numerous anthropogenically induced environmental changes. Top among these is the eutrophication of freshwaters from agriculture, development and land-use change. Here we use the genetic information recorded in lake sediments of two lakes experiencing contrasting histories of land-use change to study the evolution of local adaptation in the lakes’ *Daphnia pulicaria* populations. Utilizing nextRAD derived Single Nucleotide Polymorphisms (SNPs), we studied the evolutionary trajectories of *Daphnia pulicaria* in both lakes. Using gene-environment correlations and F_st_ tests for selection we found SNPs that appear to be under selection in both lakes. Specifically, we found more outlier SNPs in the highly impacted lake using Fst-based tests for selection. Conversely, gene-environment tests revealed the reverse pattern. We discuss numerous facets of experimental design that must be considered when using resurrection ecology to study local adaptation and critically evaluate how they may have impacted the results of this investigation.

**Lay Summary:** Resurrection ecology, the resuscitation or hatching of decades or centuries old dormant eggs, seeds or cysts provides the opportunity to study evolution in action. Here, we use resurrection ecology paired with Single Nucleotide Polymorphism (SNP) genotyping to study the evolutionary responses of two populations of *Daphnia pulicaria* to contrasting changes in the nutrient dynamics of their respective lakes. In the lake with more drastic changes in nutrient pollution, we find a stronger shift in allele frequencies through time and at a larger number of affected genomic positions compared with the environmentally more stable lake. However, Bayesian gene-environment correlations were stronger in the more stable lake reflecting higher power to detect correlations among allele frequency change and paleo-environmental variables in this location. Our results suggest that numerous factors might impact the ability to use different methodologies to detect local adaptation over time using resurrection ecology.

## Introduction

In his 1859 novel, *A Tale of Two Cities*, Charles Dickens compares the cities of London and Paris in the late 18th century. Juxtaposition is the central literary device Dickens uses in his novel. While bi-directional comparison is useful in literature, it can also be illustrative in biology. Specifically, in evolutionary biology, the availability of two or more populations experiencing contrasting selection regimes provides the basis for demonstrating local adaptation (Kawecki and Ebert, 2004). The occurrence of local adaptation and spatial structuring of habitats provides the basis for the diversification of lineages and the origin of biodiversity by the exploitation of ecological opportunity (Rainey and Travisano, 1998).

Studies of local adaptation typically compare two or more contemporaneous populations arrayed in space across a measurable environmental gradient. Differences in allele frequencies (e.g., F_st_) between populations sampled from divergent environments can be used to pinpoint the genes responsible for local adaptation by finding those that deviate from the expectation of neutrality (Lewontin and Krakauer, 1973). More recently, studies of local adaptation have also sought to find correlations between genetic markers and environmental variables to better understand the connection between measures of genetic distance and the relation these have with the environment (Günther and Coop, 2013; Rellstab et al., 2015). Regardless of the approach used, these methods universally rely on the assumption that studies of local adaptation can substitute spatial distance and its correlation with environmental distance as a proxy for the operational timeframe of evolutionary dynamics in the system. This paradigm, called “space-for-time substitutions,” is not without merit, however data interpretation may be confounded by other factors such as non-equilibrium population histories, historical contingencies and genetic drift or mutation that influence the genetics of the study populations (Lovell et al., 2023; Vermeij, 2006). An approach that may ameliorate these factors would be to study a population *in situ* during evolution by sampling individuals across different points in time.

The availability of extended allele frequency trajectories together with contemporaneous habitat conditions affords a unique opportunity to observe evolutionary change through time. When combined together, there is the potential to develop insights in the evolution of local adaptation by the identification of underlying genetic variants responsible for phenotypic change (Wersebe and Weider, 2023). Such inference may be further heightened when considered within the context of larger spatial and temporal heterogeneity. Specifically, comparing one population experiencing significant ecological disruption to one experiencing relatively stable conditions may provide the ability to specifically account for patterns of local versus global adaptive variation. Such data can be collected for *Daphnia*, a keystone microcrustacean grazer in freshwater environments, either by direct sequencing of dormant eggs deposited in the layered sediment of lakes (Lack et al., 2018; O’Grady et al., 2022) or by sequencing the extracted DNA of ‘resurrected’ *Daphnia* isolates (Kerfoot et al., 1999; Kerfoot and Weider, 2004). Here we use the second approach to obtain genomic information on historic and extant populations of *Daphnia pulicaria* in two lakes located in Minnesota, USA.

*Daphnia* are considered keystone species in lake food webs, connecting the flow of energy from algal primary producers to higher-level consumers such as fish (Lampert, 2011). Additionally, large-bodied *Daphnia* species such as *D. pulicaria,* provision key ecosystem services including the maintenance of water clarity by reducing the standing crop of algae and supporting recreational fisheries (Walsh et al., 2016). Previous work in two lakes in Minnesota (U.S.A.), South Center (SC) Lake and Hill Lake (Frisch et al., 2014, 2017) demonstrated that since widespread Western European colonization of North America, SC Lake has experienced cultural eutrophication, transitioning to eutrophic conditions within the last 600 years with a marked shift in phosphorus conditions and lake productivity which peaked between the 1970s and 1990s (Frisch et al., 2014, 2017). In contrast, Hill Lake has not experienced strong shifts in nutrient loading, and its environmental conditions have remained relatively stable across the past ∼250 years (Frisch et al., 2017).

Extending the opening metaphor, Dickens, in a *Tale of Two Cities* highlighted the divergence of revolutionary Paris to 18th century London, a hallmark of conservative stability. Here, we utilize RADseq derived SNPs to uncover the genetic tale of local adaptation of two populations of *Daphnia pulicaria* experiencing contrasting environmental regimes. Specifically, we hypothesized that: (1) more genetic markers experience extreme changes in allele frequency over time in the population under stronger anthropogenic pressure in SC Lake compared to that of the relatively more stable Hill Lake because more drastic changes in environmental conditions would necessarily spur local adaptation; (2) the lake populations have diverging evolutionary trajectories (different gene families are impacted) due to different selection regimes; (3) genomic adaptation in each lake can be attributed to environmental history of a selection of proxies reconstructed from lake sediments.

## Methods

### Study area and sampling

Sediment cores were extracted from South Center (SC) Lake (45°22.645′ N, 92°49.215′ W) and Hill Lake (47°1.1520N, 93°5.9000W), Minnesota, USA in July 2010 and 2011. For a detailed methodology of coring and radiometric dating see Frisch et al. (2014, 2017). For this study, we isolated a total of 95 clones from lake water in 2010/2011 or resurrected them from sediment layers. In SC Lake they represent five temporal subpopulations of the years 2011, 2007, 2001, 1977 (10 clones each) and 1530 (date is midpoint between two time periods that were merged - two clones from 1648 and two clones from 1418), and six temporal subpopulations in Hill Lake of the years 2010 (10 clones), 2007 (12 clones), 2002, 1997 (10 clones each),1990 (6 clones) and 1974 (date is midpoint between three time periods that were merged - one clone each from 1983, 1976 and 1962). We hatched all clones from dormant eggs collected in the sediment as described in Frisch et al. (2014) except those of the years 2010 and 2011, which were directly sampled from the lake population.

### Molecular

#### DNA extraction and Radseq sequencing

For each of the 95 clones, genomic DNA was extracted from 10 adult *Daphnia pulicaria* (Forbes 1893) raised as isoclonal cultures, using a modified CTAB protocol (Doyle & Doyle, 1987). Genomic DNA was converted into nextRAD genotyping-by-sequencing libraries (SNPsaurus LLC) as in Russello et al. (2015). Briefly, genomic DNA was first fragmented with Nextera reagent (Illumina, Inc), which also ligates short adapter sequences to the ends of the fragments. The Nextera reaction was scaled for fragmenting 2 ng of genomic DNA, although 2.25 ng of genomic DNA was used for input. Fragmented DNA was then amplified for 25 cycles at 73 °C, with one of the primers matching the adapter and extending seven nucleotides into the genomic DNA with the selective sequence GTATAGG. Thus, only fragments starting with a sequence that can be hybridized by the selective sequence of the primer were efficiently amplified. The nextRAD libraries were sequenced on one lane of an Illumina HiSeq 2000 to generate single-end sequencing reads (University of Oregon, USA).

#### Bioinformatic processing

We first de-multiplexed and quality-filtered the Illumina sequencing reads using the process_radtags command in the STACKS program (Catchen et al., 2013). We aligned the de-multiplexed RAD sequencing libraries to the *Daphnia pulicaria* reference genome (Wersebe et al. 2022; RefSeq: GCF_021234035.1) using the BWA mem algorithm (Li and Durbin, 2009).The resulting alignments were piped to SAMtools (Li et al., 2009) to mark PCR duplicates, sort the alignments and write BAM files. Next, we passed the BAM files to BCFtools to call SNPs using the mpileup and call subcommands (Danecek et al., 2021). We called SNPs in the data set in a population-specific manner (e.g., Hill and SC). We filtered both resulting VCF files to a set of high confidence bi-allelic SNPs using BCFtools, excluding loci with a read depth <10, phred quality < 30, mapping quality < 10 and a missing rate >0.4.

#### Population Structure

For each lake, we grouped samples into ‘temporal subpopulations’ based on the sediment depth from which they were hatched. We determined the approximate age of each subpopulation using the radiometric dating model (see above). We computed PCAs to evaluate the structure of temporal populations, separately for each lake. To produce the SNP set for this analysis, we pruned the set of high quality bi-allelic SNPs available for each lake for linkage disequilibrium using PLINK (Chang et al., 2015). PCAs were computed with the R package SNPrelate (Zheng et al., 2012), and visualized with ggplot2 version 3.4.2 (Wickham, 2016).

We further explored population genetic structure with a discriminant analysis of principle components (DAPC) with the R package adegenet version 2.1.10 (Jombart, 2008; Jombart et al., 2010). We used the function ‘xvalDAPC()’ to estimate how many axes should be retained for the final discriminant analysis (South Center Lake: 15, Hill Lake:10).

PCA and DAPC were computed on the R platform version 4.2.2 (R Core Team 2022).

#### Outlier Analysis

We used two methods per lake to identify outlier SNPs present in the data set. The first method, relied on the framework proposed by Wersebe and Weider (2023), which searches for SNPs with F_st_ outside the neutral expectations determined by demography. As mentioned above, for each lake, we grouped samples into ‘temporal subpopulations’ based on the sediment depth from which they were hatched. We determined the approximate age of each subpopulation using the radiometric dating model (see above) and we calculated the number of generations between each subpopulation using a fixed 5 generations per year. For both lakes, the clones isolated from the water column (e.g., lake-clones) represented generation 0 with generation estimates for older subpopulations relative to this benchmark. Next, we pruned the set of high quality bi-allelic SNPs available for each lake for linkage disequilibrium using PLINK (Chang et al. 2015). Using these LD-pruned SNPs, we estimated two-dimensional folded site frequency spectra (SFS) for each pairwise subpopulation comparison using the program easySFS (GitHub: https://github.com/isaacovercast/easySFS) formatted for the program FastSimCoal2. We fit a simple demographic scenario for each lake, where we estimated the effective population size (N_e_), a growth rate parameter and historical sampling using the observed SFS in the maximum-likelihood framework provided in FastSimCoal2. Briefly, we estimated the best fitting parameters by launching 100 independent simulations using 1-million coalescent simulations and 40 Brent maximization cycles. We found the best fitting parameters by extracting the simulation run with the highest estimated likelihood from the 100 simulation runs. We conducted this process for both Hill and South Center separately. From South Center (SC) Lake we excluded the two oldest sub-populations (60-64 cm & 52-56 cm) from this analysis because the 60-64-3X clone had low overall coverage resulting in few recovered loci and the inaccuracy of estimating F_st_ from small sample sizes (i.e., two clones). A similar approach was used for Hill as well, where singleton clones sampled from the oldest layers were removed.

Next, we used FastSimCoal2’s coalescent simulator to generate genetic markers under the inferred demographic parameters using 100 independent simulations. The SNPs simulated under the inferred demographic model were used to calculate empirical p-values for the SNPs observed in the actual populations. For each set of simulated SNPs, we converted the native FastSimCoal2 format (arlequin) to VCF using PGDSpider (Lischer and Excoffier, 2012). We calculated an estimate of site-wise F_st_ in R using the package heirfstat using the function basic.stats (Goudet, 2005). The simulated F_st_ values were used to construct a distribution for F_st_-values expected under neutral demography. We tested each of the estimated F_st_ values for the observed SNPs and determined the presence of outliers by extracting SNPs with false discovery rate (FDR) corrected F_st_ p-values above a p = 0.05 threshold.

In addition to the simulation-based approach, we used as a second approach the outlier detection program Bayenv2 (Günther & Coop 2013) to detect SNPs potentially correlated with environmental variables inferred from the sediment cores. This program requires three input matrices that were constructed according to the Bayenv2 manual: an environmental matrix, a SNPs matrix and a covariance matrix. The environmental matrix for both lakes contained proxies for lake productivity that were estimated by paleolimnological methods using ^210^Pb-dated sediment (for details see Frisch et al. 2017): sediment age, accumulation rates (flux) of organic carbon (OC), calcium carbonates (CaCO_3_), and ortho-phosphorus (P) for both SC and Hill lakes, estimated for the 6 and 5 subpopulations in SC and Hill, respectively.

The covariance matrix implemented by Bayenv2 was estimated from a set of putatively neutral intergenic SNPs to account for changes in allele frequencies related to population history and sampling bias. To fit the required population covariance matrix, we extracted all intergenic SNPs present in the lake-specific and jointly called VCF files using the R packages GenomicRanges 1.50.1 and GenomicFeatures 1.50.4 (Lawrence et al., 2013). Briefly, we intersected the *Daphnia pulicaria* RefSeq gff3 (NCBI RefSeq assembly GCF_021234035.1) containing only gene annotations with the above described VCF file of high confidence SNPs to extract intergenic SNPs. This SNP set was then filtered using the R package SNPrelate to obtain biallelic LD pruned SNPs with a minimum allele frequency of 0.05, missing rate of 0.25, yielding 3509 SNPs for SC Lake and 2636 SNPs for Hill Lake. The covariance matrix was estimated using 500,000 iterations. SNP sets for each lake were tested against the respective covariance matrix to test their correlation with the environmental matrix with 500,000 iterations and was repeated five times. We set the criteria for SNPs significantly correlated with one of the tested environmental factors as at least a median Bayes factor across the five runs of 2.0 or higher, or at least two runs with Bayes factors of 2.0 or higher.

#### Power assessment and false positive rates for Bayenv2 analysis

The two sets of intergenic SNPs (one per lake) as well as the covariance matrices estimated from them were used to conduct simulations for assessing power and false positive rates of Bayenv2 under different criteria. Out of all intergenic SNPs, we randomly picked 1000 SNPs to create Bayenv2 input files with a simulated environmental effect on their allele frequency. For each SNP and each population *i*, the empirical allele frequency was calculated by dividing the total number of chromosomes carrying the allele *n_i_* by the total number of genotyped chromosomes for population and site *N_i_*. Using the normalized environmental variable *Y* (mean 0, standard deviation 1) as well as an effect strength *β*, the simulated allele frequency *f_i_* was calculated by adding a linear environmental effect *β_i_* · *Y_i_* to the the empirical point estimate 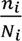 of the allele frequency. If the result fell below 0 or was greater than 1, the simulated frequency was set to 0.0 or 1.0, respectively. The calculation of *f_i_* is described in Equation (1) below.

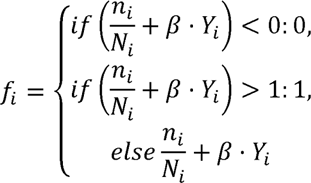

Equation (1):

Allele counts for the Bayenv2 input files were then calculated by rounding *f_i_·N_i_* to the nearest integer. The remaining chromosomes for each population were assumed to carry the alternative allele.

This resulted in 1000 SNPs per combination of environmental variable, effect strength β and test case. For each simulated SNP, Bayenv2 was then used to estimate Bayes factors with five different random seeds using the empirical covariance matrix. Two criteria were considered for the identification of outlier SNPs: a median Bayes factor across the five runs of 2.5 or higher, or at least three runs with Bayes factors of 2.0 or higher (Fig. S1 and S3 for SC Lake and Hill Lake, respectively). The estimates for the power to detect such outliers were then calculated as the proportion of the simulated SNPs exceeding those criteria. Similarly, the false positive rate was estimated as the proportion of the original intergenic (i.e. presumably neutral) SNPs exceeding these criteria across five independent runs of Bayenv2 (Fig. S2 and S4 for SC Lake and Hill Lake, respectively).

#### Functional Enrichment

We sought to understand the functional context of regions hosting outlier SNPs identified with both outlier detection methods. We accomplished this by identifying enriched Gene Ontology (GO) terms for the genes related to outlier SNPs. We annotated the effects of all SNPs identified in both lakes using the program Variant Effect Predictor (VEP; (McLaren et al., 2016)), which provides both an effect of a SNP and the genes it is plausibly related to. Using this context, we were able to annotate which genes were related to the SNPs identified as outliers using the RefSeq *Daphnia pulicaria* annotation. After extracting the genes related to outlier SNPs, we annotated these genes with their PantherDB generic mappings. Using the PantherDB generic mappings for outlier-associated genes, we identified enriched GO terms via the PantherDB webtool using *Daphnia pulex* genes as a reference (Mi et al., 2021).

## Results

Our sequencing efforts produced 95 single-end NextRAD libraries. In total, we sequenced 44 distinct clones from the South Center (SC) population and called 9505 high confidence bi-allelic SNPs from the five temporal subpopulations of this lake. We sequenced 51 samples from the six Hill temporal subpopulations, which yielded 6939 high confidence bi-allelic SNPs in the population.

### Population genetic structure across time

Using LD pruned SNP sets for both lakes, we computed graphic representations of PCAs and DAPCs (Fig. 1A-1C). For both lakes, the PCAs show a minor population structure with a variance between 5 and 7% on PC1, and around 5% on PC2 (Fig. 1, top panels). In SC Lake, much of this variation can be explained by the 1970 subpopulation that forms a separate cluster.

**Figure 1:**
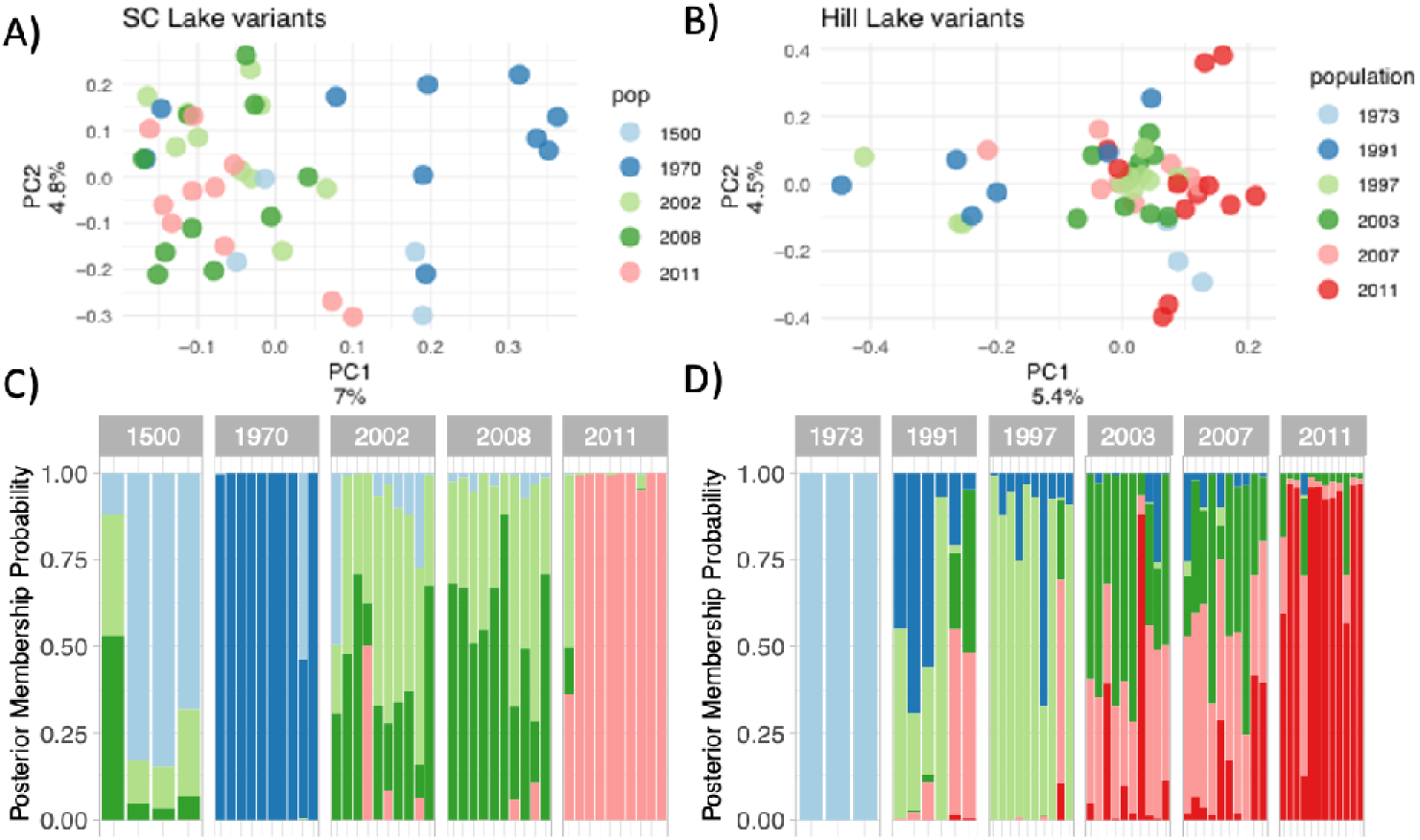
Population Genetic Structure. A) South Center (SC) Lake individual PCA Biplot. B) Hill Lake individual PCA Biplot. C) Discriminant Analysis of Principle Components (DAPC) posterior membership probabilities for SC Lake individuals. D) DAPC posterior membership probabilities for Hill Lake individuals.

In Hill Lake, none of the populations cluster distinctly from the others, and the variance is related to individuals from several temporal subpopulations. The discriminant analysis for SC resulted in distinct groupings of the subpopulations, assigning most of the individuals clearly to the sediment depth from which they were resurrected. The results of the DAPC for Hill were less distinct with mixed assignments of individuals for several subpopulations, indicating less differentiation between the Hill temporal subpopulations.

### FSC Outliers

We estimated F_st_ for all sites recovered from the populations of each lake. We compared these against our simulated estimates for expected neutral F_st_ and estimated empirical p-values (Fig. 2- L & R). For the SC Lake population, observed F_st_ ranged from -0.1054 to a high of 0.7703 (Fig. 3-L). The mean F_st_ across all sites was low overall at 0.032. In total, we identified 122 outlier SNPs with FDR adjusted p-values above a significance threshold of 0.05, these SNPs all had F_st_ estimates at or above 0.2547.

**Figure 2:**
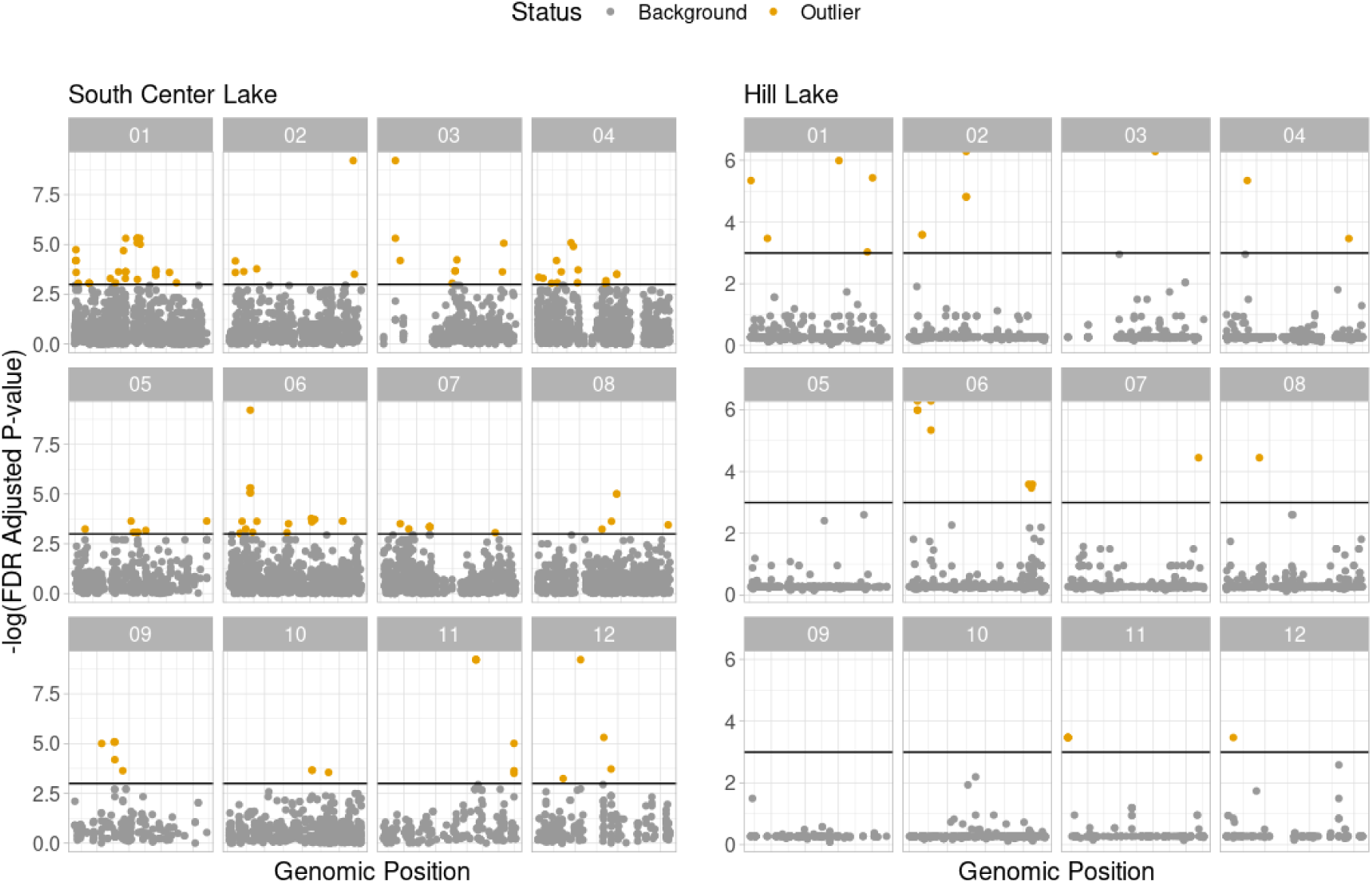
F_st_ outliers. Left) F_st_ outliers identified in South Center (SC) Lake. Right) F_st_ outliers identified in Hill Lake. Grey points are SNPs without significant F_st_ values based on simulation Gold points are SNPs with significant F_st_ values based on simulations. Vertical bars denote p = 0.05 cutoff.

**Figure 3).**
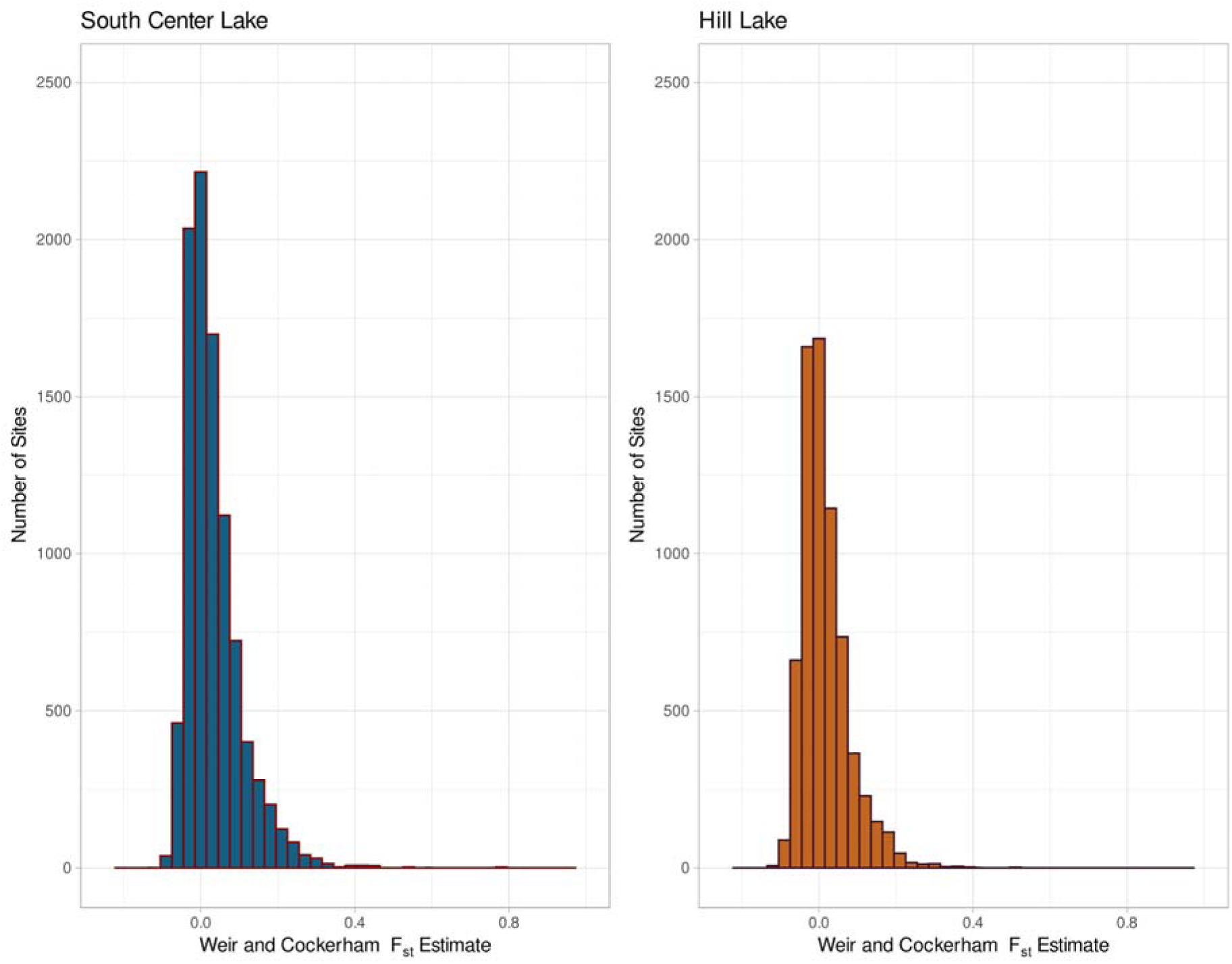
Observed F_st_ distbution. Left) South Center (SC) Lake. Several sites reach F_st_ near 0.77. Right) Hill Lake. F_st_ reaches a high of 0.55.

For the Hill Lake population, the dynamics of outliers diverged significantly from those observed in the SC population. Observed site-wise estimates of F_st_ ranged from a low of -0.1219 to a high of 0.519 (Fig. 3-R). The mean F_st_ across the subpopulations was also relatively low at 0.016. In total, however, we observed just 29 outlier loci in the entire data set, all of which had estimated F_st_ at or above 0.2839.

### Functional Enrichment

To better understand the genomic context of outlier regions, we extracted the genes related to high F_st_ SNPs and annotated them with Gene Ontology (GO) terms.

#### F_st_ Outliers

We extracted a total of 120 unique genes that were related to high F_st_ SNPs identified in the SC population. We were able to annotate 118 of these genes with functional family information while searching against the Panther database. This list of genes yielded 32 GO terms for Molecular Function that appeared enriched after using a Fisher exact test against the *Daphnia pulex* gene list available on PantherDB webtool and correcting for FDR. Enriched GO terms were related to several molecular functions; however, transmembrane transport proteins for both organic molecules and ions and molecule-specific binding made up many of the enriched terms. Meanwhile, for the Hill Lake population, we only identified 24 genes related to outlier SNPs. This yielded no enriched terms for molecular function, and the gene list contained only two genes that were found in South Center.

#### Bayenv2 Outliers

We identified only five outlier loci in the SC population, which were related to six genes. This relatively small number reflects the low power of Bayenv2 to detect environmental outliers in South Center (Fig. S1 and S3), likely originating from the strong correlation between the main axis of population structure (Fig 1A) and the environmental variables which caused Bayenv2 to over-correct for this structure. The six genes resulted in no enriched GO terms for molecular function. The Hill population had loci correlated with age, CaCO_3_, Organic Carbon (OrgC) and phosphorus (P) ranging from a high of 43 loci correlated with OrgC content to 5 loci correlated with P (Table S1). The small number of genes related to outlier SNPs from age, P, and CaCO_3_ did not result in any enriched GO terms. However, SNPs associated with organic carbon were related to a total of 58 genes. This gene list resulted in 10 enriched GO terms which were related to several transcription factors including binding and regulation.

**Figure 4).**
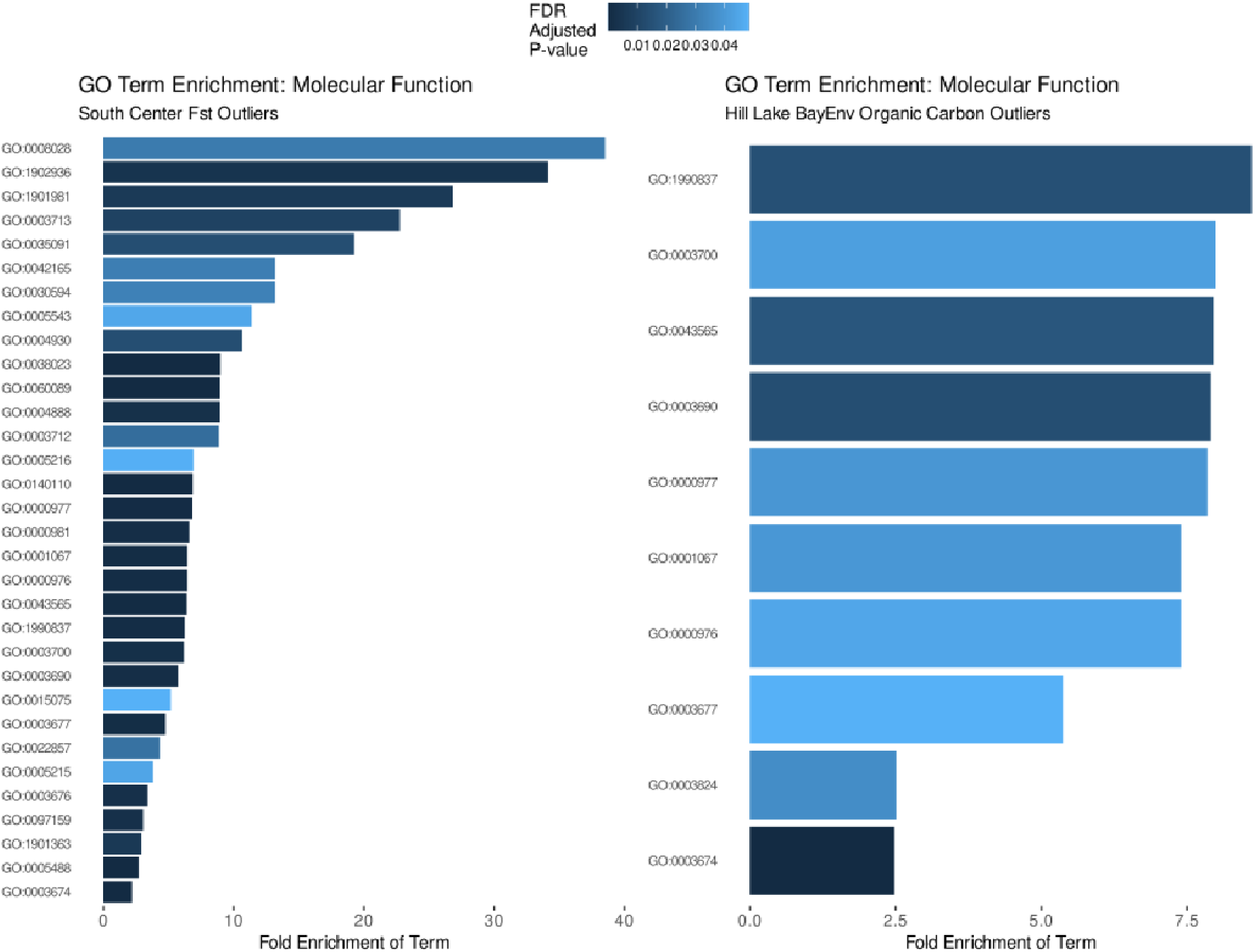
Outlier SNP GO term Enrichment. Left) PANTHERDB GO term for molecular function enriched among F_st_ outlier SNPs identified in SC Lake. Right) PANTHERDB GO term for molecular function enriched among Bayenv outlier SNPs identified in Hill Lake. The GO terms corresponding to IDs here are present in tables S2 (L) and S3 (R).

## Discussion

Our data support our first hypothesis - that significant anthropogenic changes within a lakes’ watershed would result in more genetic markers experiencing dramatic changes in allele frequency. We found only limited support for our other two hypotheses that lake populations would have diverging evolutionary trajectories related to different selection regimes and that genomic adaptation can be attributed to selected environmental proxies. In many cases, similar GO terms for molecular function were enriched in both lakes and few SNPs had measurable gene-environment correlations.

### Human impacts leave measurable signatures in the genome

The identification of F_st_ outliers revealed more than four times the number of outliers in SC Lake compared to those observed in Hill Lake (i.e., 122 sites vs 29 sites). This supports our hypothesis that more sites would be under selection in the *Daphnia* population of SC Lake, resulting from the dramatic environmental changes in this lake. The SC lake watershed is dominated largely by pastureland and row-crop agriculture encompassing 38% and 23% of the watershed area, respectively (MNPCA, 2009a). As noted previously (Frisch et al. 2014, 2017) this has created a set of novel environmental conditions in SC Lake, particularly because of nutrient loading (N & P) from agricultural run-off. The watershed of Hill Lake, by contrast, is largely dominated by forest covering approximately 75% of the watershed whereas 14% of the land is used by agriculture resulting in mesotrophic conditions (MNPCA, 2009b). This provides further evidence of the enormous adaptive capacity of *Daphnia* populations at the genomic level. Previous studies have shown that resurrected individuals from the SC population have evolved phenotypic plasticity in their phosphorus (P) physiology that allows them to regulate the retention of P according to availability in the environment (Frisch et al., 2014; Chowdhury and Jeyasingh, 2016; Frisch et al., 2020). Other resurrection studies have shown an adaptive capacity to a variety of environmental challenges including salt pollution (Wersebe and Weider 2023) or rising temperatures (Geerts et al 2015). This suggests that *Daphnia* populations will continue to support aquatic food webs and maintain the ecosystem services they provide (Walsh et al. 2016) as long as the pace or strength of environmental stress does not overwhelm their adaptive capacity.

### Many SNPs, no coherent genes

Our results from the SC population F_st_-tests are congruent with other recent studies which have used resurrection ecology paired with genome-wide markers to study local adaptation. Both Chaturvedi et al. (2021) and Wersebe and Weider (2023) found that a large number of SNPs may rapidly change in allele frequency in *Daphnia* populations experiencing novel environmental conditions. However, these studies employed whole-genome sequencing to identify SNPs segregating in the study populations over time. For example, Wersebe and Weider (2023) were able to use their data set to identify genes and mutations with known functions that were related to the physiology of the trait (i.e., salinity tolerance) they suspected to be under selection in the study population. In contrast, Chaturvedi et al. (2021) did not attempt to identify the genes or functional implications of the SNPs detected in their study.

Here, our effort to identify genes associated with the F_st_-outlier SNPs did not reveal distinct physiological functions that might be plausibly related to eutrophication. For instance Weider et al. (1997), proposed that allozyme variation at the phosphoglucose isomerase (PGI, EC 5.3.1.9) gene was related to micro-evolutionary changes associated with eutrophication in the Lake Constance (Bodensee, Germany) *Daphnia* population. Further experimental work has suggested that PGI-genotype may indeed play a role in the competitive ability of *Daphnia* under contrasting phosphorus supply (Jeyasingh et al., 2009). Despite the lack of a single locus of large effect with a clear physiological connection, we did find similar sets of genes enriched as those detected by Muñoz et al. (2016). These authors analyzed SNPs detected with genotype-by-sequencing from several *Daphnia pulicaria* populations in Minnesota including SC and Hill Lakes.

Many of the outlier SNPs detected here were enriched in GO terms related to regulation of transcription, but not immediately connected to nutrient physiology. The lack of a coherent list of genes may be related to several factors. We employed a variant of RADseq which is known as a reduced representation method. RAD genotyping only samples loci in the genome near enzyme cut sites and as a result, only samples a small portion of the DNA (Andrews et al., 2016). Some have questioned this method for studying local adaptation (Lowry et al., 2016), because the RAD approach lacks sufficient marker density to sample sites in tight linkage disequilibrium with a selected site. As such, we may have lost some power to detect selection in the genome.

Furthermore, since coverage is restricted to RAD tags, we were unable to search regions surrounding outliers for SNPs within genes causing missense mutations or premature stop codons. Such non-neutral variation is the target of selection, which might narrow a list of nearby candidate genes when examined more closely.

### Paleo-environmental correlations with allele frequency change

While the F_st_ analysis supported our hypothesis, the Bayenv analysis did not produce congruent results. This analysis detected most outlier SNPs in the Hill Lake population, while a much lower number was found in the SC Lake population. As demonstrated by the Bayenv simulations (Fig. S1,S2,S3, S4), the power of detecting outlier loci in SC Lake was very low due to the strong correlation between variation in the SNP data and environmental variables. While Bayenv2 is designed to avoid false positives due to background population structure, it might over-correct if the population structure is highly correlated with the environmental variables reducing the chances of finding loci involved in environmental adaptation. This strong correlation between genomic and environmental variation across time might explain why more SNPs with high F_st_ could be detected when applying an F_st_ approach without considering environmental data.

For Hill Lake, this correlation was less pronounced, allowing a greater power of detecting outliers associated with environmental variables. In this lake, the only environmental factor with many correlated outliers was organic carbon (OrgC) flux. This OrgC paleo-environmental record tracks the burial of organic matter in lake sediments (Tranvik et al., 2009). Studies of Minnesota Lakes have revealed that one of the primary controls of lake OrgC burial is a change in land use within a lake’s watershed (Anderson et al., 2013). As noted previously, land use is strikingly different between Hill and South Center, with the most dramatic changes occurring within South Center Lake’s watershed. These results remain difficult to explain in the context of the data we collected.

There are several factors that might impact studies such as ours, many of them inherent with difficulties related to the design of resurrection ecology studies. Our sampling of individuals throughout the core relies solely on *Daphnia pulicaria* lineages that hatched and survived in laboratory culture. This may have resulted in a non-representative sampling consisting of those genotypes that survived extended diapause and laboratory conditions. This produces a bias termed the “invisible fraction”, which arises when strong correlations exist between propagule survival and traits of interest for a given genotype (Weis, 2018). However, such biases are not directly quantifiable without knowledge of the ancestral relationships of hatched propagules. In previous studies (Frisch et al. 2014, 2017), the sampling design would have alleviated the impact of the invisible fraction because eggs were recovered from the sediments and genotyped directly. These studies demonstrated compelling evidence of gene-environment correlations, which we did not recover despite the larger number of markers employed in the present study. The larger number of sampled haplotypes in these studies may have improved the ability to detect allele frequency shifts which underly the reported gene-environment correlations.

Frisch et al. (2014) also observed phenotypic changes in the SC Lake population after profiling some of the clones included in this present study for phosphorus use efficiency and growth rate (e.g., fitness) under contrasting nutrient conditions. These observations may be related to transcriptomic and/or epigenomic changes across time rather than selection on genetic variation in the genes underlying physiological traits. Further, Frisch et al. (2020) mapped the transcriptional networks associated with P-use traits in the SC population. While many transcriptional responses were shared between the ancient and modern clones profiled, a small number showed novelty in the modern clones under P-limitation. Such a mechanism might explain in part why here, we uncovered several genes related to regulation and translation rather than to nutrient physiology. This highlights the importance of designing studies that incorporate the full complement of functional genomic interrogation including full genome sequences, transcriptomic and epigenomic profiles and common garden experiments to understand the genetic basis of phenotypic change in wild populations. Such a comprehensive and multi-faceted study using the resurrection approach has yet to be conducted.

### Challenges and Considerations of the “resurrection” approach

Many studies utilizing the resurrection approach consider the evolutionary trajectory of a single lake and its target population (see Weider et al. 2018 for an overall review). While this approach may be powerful, results from a single population are inherently phenomenological without replication or control. The addition of a “control” population such Hill Lake in our study allows us to demonstrate a concerted change in South Center Lake. However, there are several challenges and considerations that any chosen design introduces that complicate a direct comparison. Specifically, it is important to have sufficient sample sizes across time periods and lakes to make temporal comparisons meaningful. One of the primary differences between our analytical approaches was whether the oldest clones recovered were included in the results.

Since it is difficult to accurately determine allele frequencies from such small sample sizes, we opted not to include these in the F_st_-based analysis. However, they were included in the Bayenv analysis. This may explain in part the contrasting results of these approaches but more importantly it also highlights the importance of acquiring a large enough sample size from each temporal subpopulation. Additionally, to allow for a meaningful comparison between two or more lakes, populations should be sampled on the same temporal scale. With the exclusion of the oldest samples from both lakes, our data sets spanned a similar timeframe (30 years in SC Lake, 20 years in Hill Lake), but the Hill Lake core was more densely sampled compared to the SC Lake core. This has unknown effects on the inference that can be gleaned from the data.

### Conclusions

The South Center and Hill populations tell a tale that spans decades to centuries. However, the genes in these populations are not easy to read. Ultimately, the *Daphnia pulicaria* populations in each lake show distinct patterns of adaptation, which coincide with many more F_st_ outliers in the SC Lake population than in Hill Lake. Most outlier SNPs are related to transcriptomic modification and regulatory genes suggesting that adaptation in these lakes is related to a complex molecular rewiring rather than one or a few major-effect genes with known physiological function. However, when conducting resurrection studies, we recommend that care and attention be paid to sample size. This can help to increase the likelihood that the analyses produce meaningful results. This is a challenge inherent in resurrection ecology because idiosyncratic deposition and preservation of eggs within sediment cores are ultimately unknowable before selecting lakes and the temporal resolution of ephippial sampling. Regardless of current limitations, the further refinement of resurrection ecological studies has the potential to provide a powerful means of examining temporal genome-environment interactions that can complement space-for-time studies (Weider et al., 2018)

## Acknowledgements

We want to thank P.D. Jeyasingh (PDJ), B.W. Culver, and R. O’Grady for assistance in the field sampling and Dan Engstrom, M.B. Edlund (MBE) and the St. Croix Watershed Research Station for assistance with earlier sediment dating. We thank E. Johnson and P. Etter (U. of Oregon) for their assistance and feedback related to the RADseq analyses. This research was supported by NSF-IOS-OEI collaborative grants #0924289 and #1256881 to LJW and #0924401 and #1256867 to PDJ. DF received additional funding from the European Union’s FP6-MOBILITY programme under the Marie Skłodowska Curie grant agreement No. 40285 and the German Research Foundation (DFG, grant FR 2582/4-1). MJW recognizes support from the Legislative-Citizen Commission on Minnesota Resources (LCMMR; 2022-272, M.L. 2022, Chp. 94, Sec. 2, Subd. 04l.). Additional support to LJW came from the Faculty Investment Program (FIP) through the U. of Oklahoma Research Council. TG was supported by a grant from Formas– a Swedish Research Council for Sustainable Development (2023-01381). Any opinions, findings, and conclusions or recommendations expressed in this material are those of the authors and do not necessarily reflect the views of the U.S. National Science Foundation or other funding agencies.

## Author Contributions

DF and PKM performed initial isolation and sequencing. DF and MJW performed analysis and wrote the first draft. TG contributed analysis and support for the Bayesian outlier detection. LJW provided funding and administration. All authors contributed to editing and reviewing the final draft.

## Data availability

All data and custom scripts will be archived on Dryad upon acceptance of this paper. Sequencing reads will be made available via NCBI’s SRA upon acceptance.

## Conflict of Interests

Authors declare no conflicts of interest.

## Notes

### Competing Interest Statement

The authors have declared no competing interest.

