## Supplement Methods and Figures for "A tale of two lakes: divergent evolutionary trajectories of two *Daphnia* populations experiencing distinct environments"

Supplementary Figures and Tables

A tale of two lakes: divergent evolutionary trajectories of two *Daphnia* populations experiencing distinct environments[[](applewebdata://4DCECA74-6091-4DAC-BE42-E56762900AE7#_blank)

Matthew J. Wersebe^1,2,*^, Torsten Günther ^3^, Philip K. Morton^2^, Lawrence J. Weider ^2^, Dagmar Frisch^2,4^

^1^ St. Croix Watershed Research Station, Science Museum of Minnesota**,** Marine-on-St. Croix, MN

^2^ Program in Ecology and Evolutionary Biology, School of Biological Sciences, University of Oklahoma, Norman, OK

^3^ Department of Organismal Biology, Uppsala University, Uppsala, Sweden

^4^ Department of Evolutionary and Integrative Ecology, Leibniz Institute of Freshwater Ecology and Inland Fisheries (IGB), Berlin, Germany

Fig. S1: Results of the BayEnv Power analysis for SC lake using two different criteria for classifying outliers. The power is calculated as the proportion of SNPs with a simulated effect of strength “beta” for the respective environmental variable.

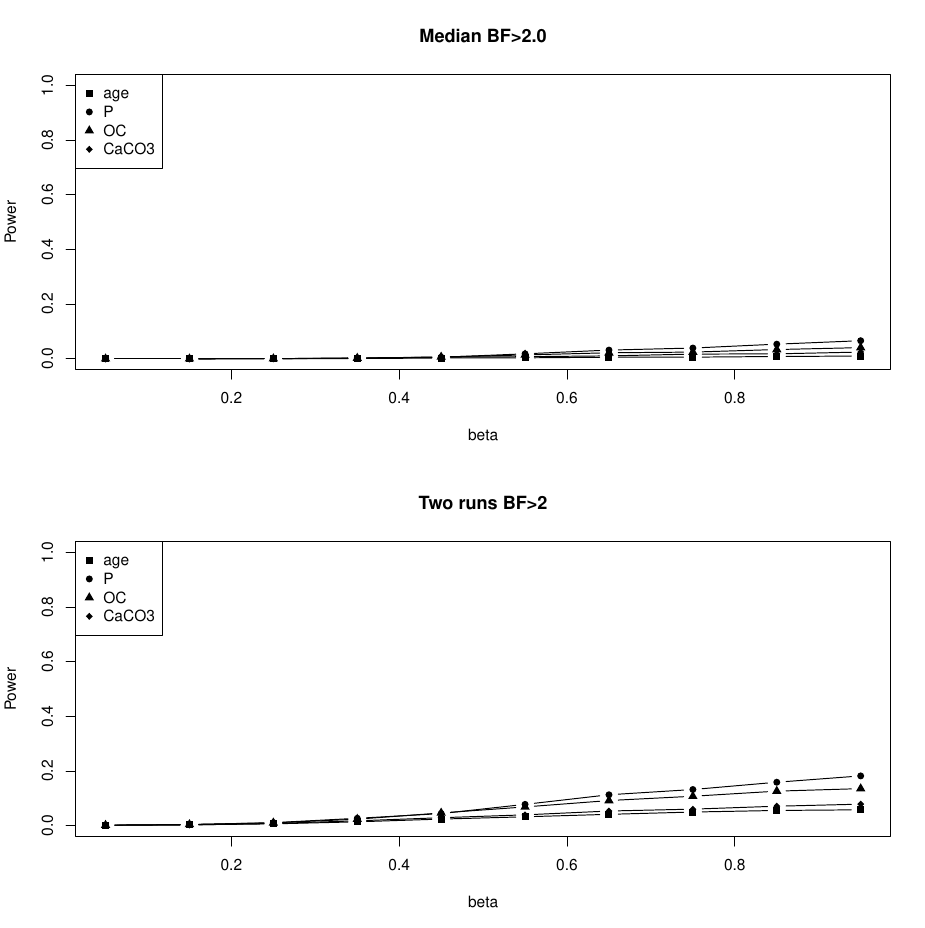

Fig. S2: Estimated false positive rate of BayEnv for SC lake. The false positive rate was estimated as the proportion of SNPs without any simulated effect that exceeded the outlier thresholds.

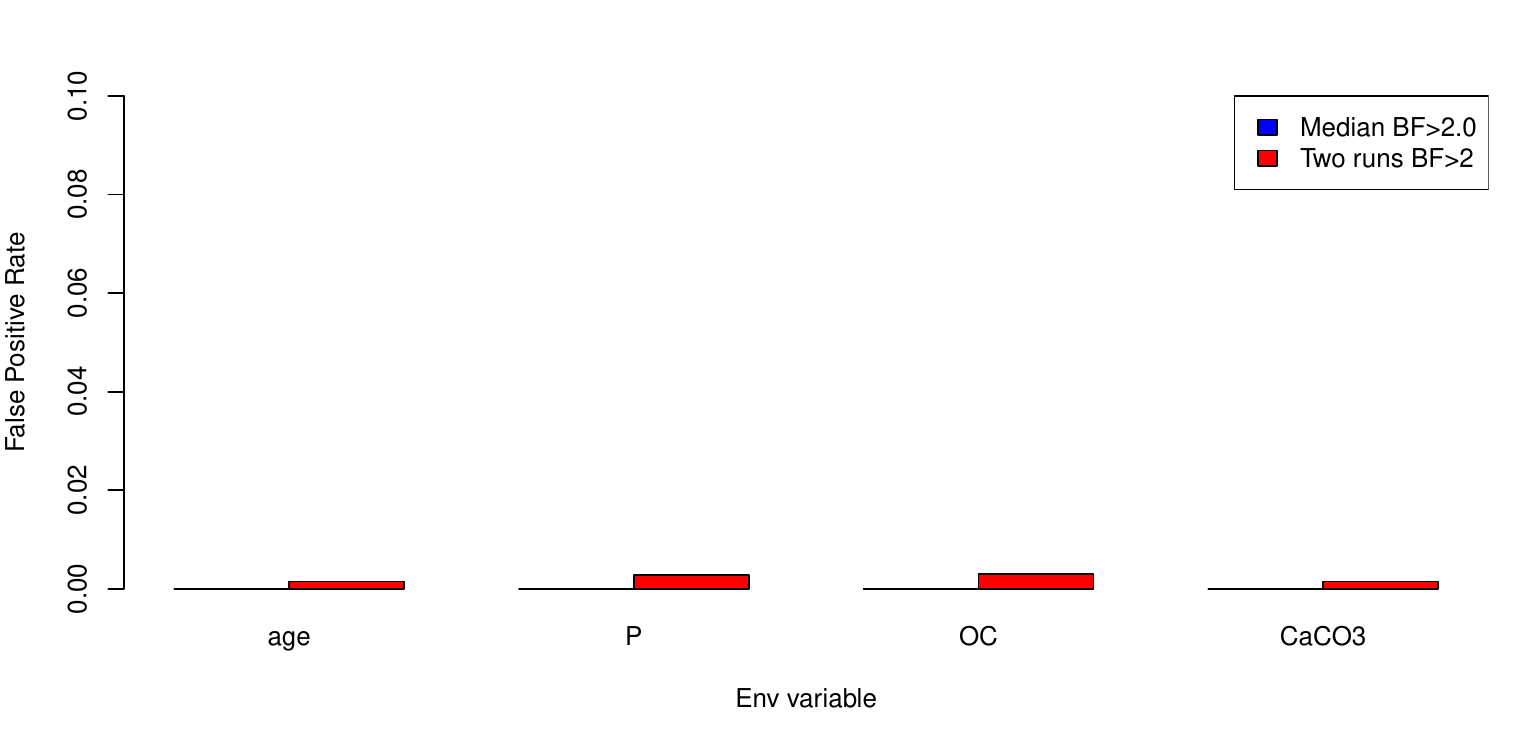

Fig. S3: Results of the BayEnv Power analysis for Hill lake using two different criteria for classifying outliers. The power is calculated as the proportion of SNPs with a simulated effect of strength “beta” for the respective environmental variable.

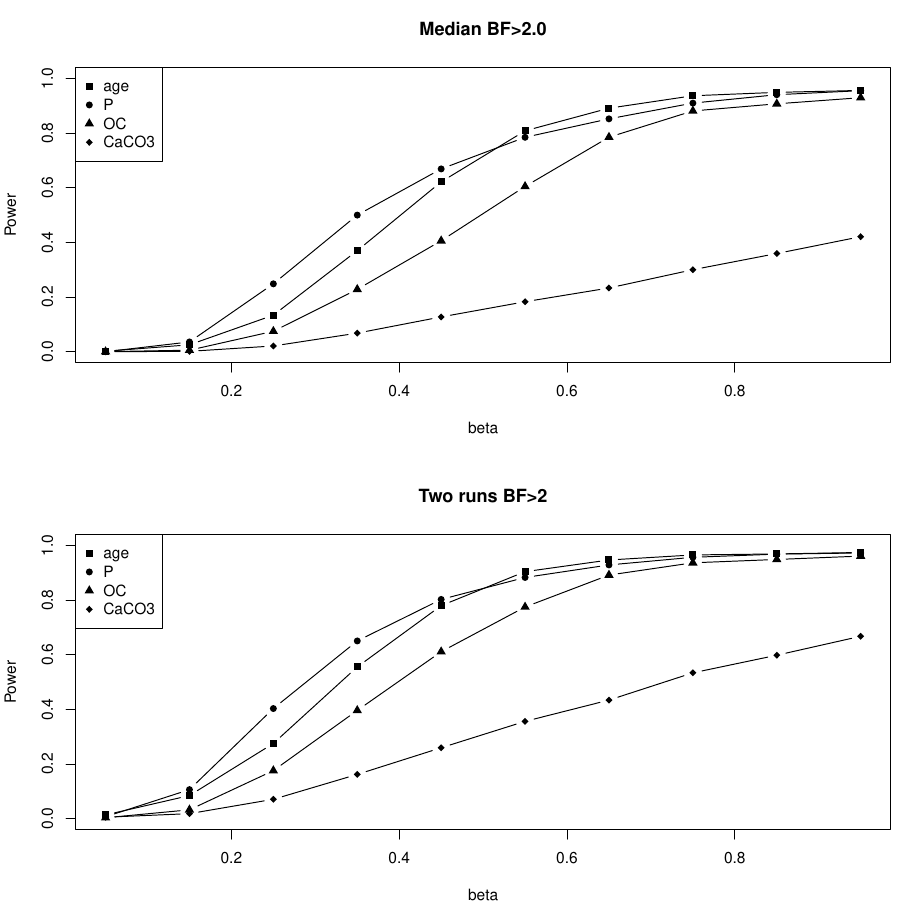

Fig. S4: Estimated false positive rate of BayEnv for Hill lake. The false positive rate was estimated as the proportion of SNPs without any simulated effect that exceeded the outlier thresholds.

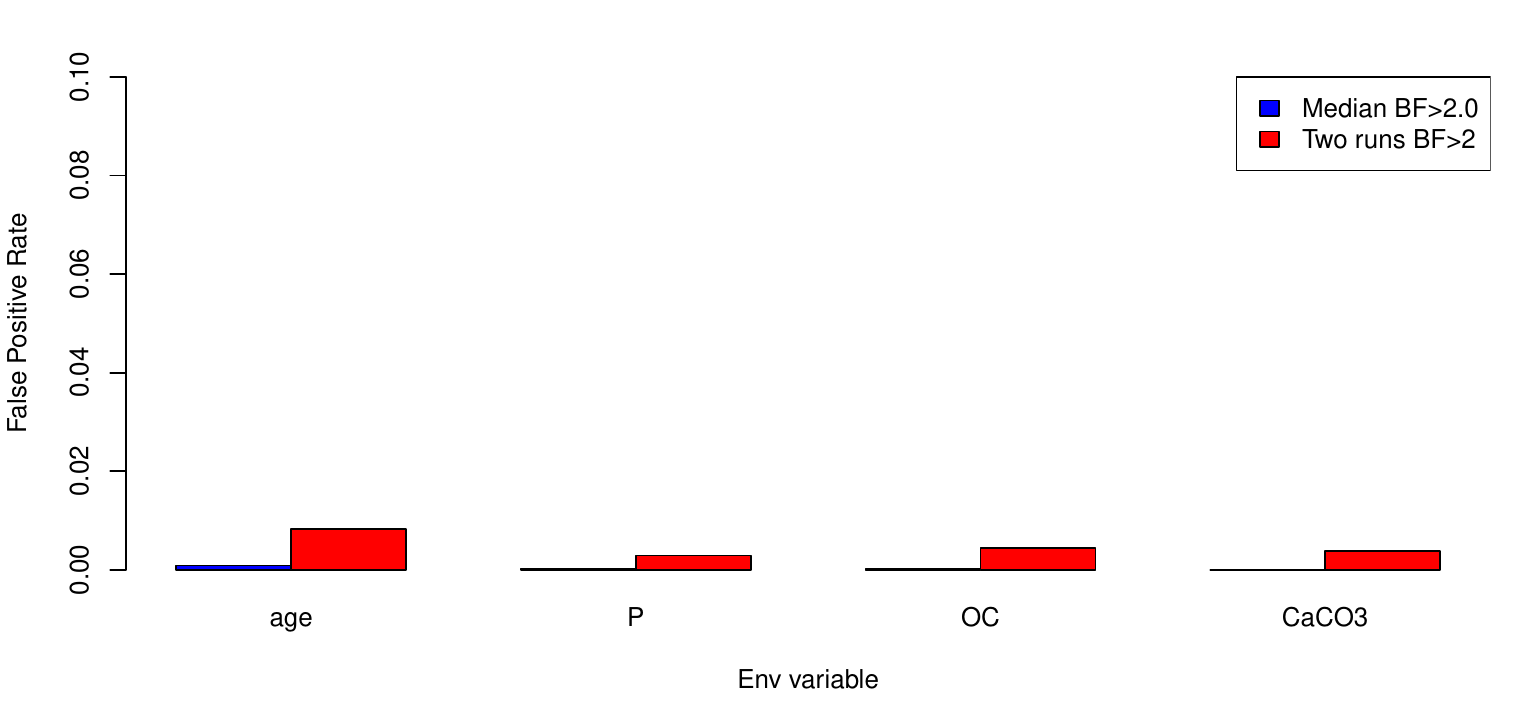

**Table S1:** Genes related to SNP outliers identified in the BayEnv analysis. Each identified gene and its ID is listed together with its product, and the lake population and environmental factor it was found to be correlated with. For details see Methods.

| **ID** | **product** | **Lake** | **env** |
| --- | --- | --- | --- |
| LOC124315508 | serine/threonine-protein kinase/endoribonuclease IRE1-like | Hill | age |
| LOC124316724 | cell surface glycoprotein 1-like | Hill | age |
| LOC124342747 | ecdysone receptor-like | Hill | age |
| LOC124316722 | endoglucanase-like | Hill | age |
| LOC124342670 | FGGY carbohydrate kinase domain-containing protein-like | Hill | age |
| LOC124342846 | kinetochore protein Spc25-like | Hill | age |
| LOC124342715 | probable tRNA N6-adenosine threonylcarbamoyltransferase%2C mitochondrial | Hill | age |
| LOC124314696 | putative ankyrin repeat protein RF_0381 | Hill | age |
| LOC124316488 | salivary glue protein Sgs-3-like isoform X1 | Hill | age |
| LOC124316525 | salivary glue protein Sgs-3-like isoform X1 | Hill | age |
| LOC124327863 | sodium/calcium exchanger 2-like isoform X1 | Hill | age |
| LOC124327863 | sodium/calcium exchanger 3-like isoform X2 | Hill | age |
| LOC124342565 | solute carrier family 12 member 3-like isoform X1 | Hill | age |
| LOC124342573 | uncharacterized protein LOC124342573 | Hill | age |
| LOC124342628 | uncharacterized protein LOC124342628 | Hill | age |
| LOC124348842 | uncharacterized protein LOC124348842 | Hill | age |
| LOC124327665 | zinc finger MYM-type protein 1-like | Hill | age |
| LOC124348859 | T-box transcription factor TBX2-like isoform X1 | SC | age |
| LOC124337068 | dynein axonemal intermediate chain 1-like | Hill | CaCO3 |
| LOC124336097 | eukaryotic translation initiation factor 4 gamma 1-like isoform X1 | Hill | CaCO3 |
| LOC124310910 | lutropin-choriogonadotropic hormone receptor-like isoform X1 | Hill | CaCO3 |
| LOC124320755 | mesoderm induction early response protein 1-like isoform X1 | Hill | CaCO3 |
| LOC124311493 | N-acetylglucosamine-6-sulfatase-like | Hill | CaCO3 |
| LOC124337148 | prolyl 4-hydroxylase subunit alpha-2-like | Hill | CaCO3 |
| LOC124311089 | uncharacterized protein LOC124311089 | Hill | CaCO3 |
| LOC124321251 | uncharacterized protein LOC124321251 isoform X1 | Hill | CaCO3 |
| LOC124336181 | uncharacterized protein LOC124336181 isoform X1 | Hill | CaCO3 |
| LOC124336247 | uncharacterized protein LOC124336247 | Hill | CaCO3 |
| LOC124336700 | uncharacterized protein LOC124336700 | Hill | CaCO3 |
| LOC124316355 | alpha-(1%2C3)-fucosyltransferase C-like | Hill | OrgC |
| LOC124348464 | alpha-(1%2C3)-fucosyltransferase C-like | Hill | OrgC |
| LOC124329274 | annexin B9-like | Hill | OrgC |
| LOC124320854 | bromodomain adjacent to zinc finger domain protein 1A-like | Hill | OrgC |
| LOC124349379 | C3 and PZP-like alpha-2-macroglobulin domain-containing protein 8 isoform X1 | Hill | OrgC |
| LOC124321228 | carotenoid isomerooxygenase-like | Hill | OrgC |
| LOC124336809 | cell division control protein 6 homolog isoform X1 | Hill | OrgC |
| LOC124342803 | cell wall protein DAN4-like | Hill | OrgC |
| LOC124337884 | chitin synthase chs-2-like isoform X1 | Hill | OrgC |
| LOC124327323 | coactosin-like protein isoform X1 | Hill | OrgC |
| LOC124335868 | cuticle protein 18.7-like | Hill | OrgC |
| LOC124335824 | cuticle protein 19-like | Hill | OrgC |
| LOC124342841 | cuticle protein 8-like | Hill | OrgC |
| LOC124326300 | enoyl-[acyl-carrier-protein] reductase%2C mitochondrial-like | Hill | OrgC |
| LOC124316498 | fibroblast growth factor receptor 2-like isoform X2 | Hill | OrgC |
| LOC124340454 | forkhead box protein G1-like | Hill | OrgC |
| LOC124316341 | G2/mitotic-specific cyclin-B3-like | Hill | OrgC |
| LOC124316352 | galactose-3-O-sulfotransferase 2-like | Hill | OrgC |
| LOC124349426 | glucose dehydrogenase [FAD%2C quinone]-like isoform X1 | Hill | OrgC |
| LOC124350679 | glutamate [NMDA] receptor subunit 1-like isoform X2 | Hill | OrgC |
| LOC124342194 | glutamate receptor ionotropic%2C delta-1-like | Hill | OrgC |
| LOC124326046 | GTPase-activating Rap/Ran-GAP domain-like protein 3 isoform X1 | Hill | OrgC |
| LOC124335866 | hatching enzyme 1.2-like | Hill | OrgC |
| LOC124335867 | hatching enzyme 1.2-like | Hill | OrgC |
| LOC124337904 | innexin inx2-like | Hill | OrgC |
| LOC124349621 | LIM homeobox transcription factor 1-alpha-like | Hill | OrgC |
| LOC124344393 | peptidyl-prolyl cis-trans isomerase FKBP8-like isoform X1 | Hill | OrgC |
| LOC124316151 | PHD finger protein 20-like isoform X1 | Hill | OrgC |
| LOC124342071 | piggyBac transposable element-derived protein 3-like | Hill | OrgC |
| LOC124326113 | poly(A) RNA polymerase gld-2 homolog A-like isoform X1 | Hill | OrgC |
| LOC124335825 | pro-resilin-like | Hill | OrgC |
| LOC124338112 | pro-resilin-like isoform X1 | Hill | OrgC |
| LOC124326408 | probable enoyl-CoA hydratase%2C mitochondrial isoform X1 | Hill | OrgC |
| LOC124310791 | protein sax-3-like isoform X1 | Hill | OrgC |
| LOC124320241 | protein trachealess-like | Hill | OrgC |
| LOC124337138 | putative cuticle collagen 80 | Hill | OrgC |
| LOC124337136 | putative cuticle collagen 91 | Hill | OrgC |
| LOC124316314 | putative zinc finger protein 833 isoform X2 | Hill | OrgC |
| LOC124336912 | rhophilin-2-like isoform X1 | Hill | OrgC |
| LOC124342785 | shematrin-like protein 1 | Hill | OrgC |
| LOC124327863 | sodium/calcium exchanger 2-like isoform X1 | Hill | OrgC |
| LOC124316136 | sporulation-specific protein 15-like | Hill | OrgC |
| LOC124342766 | transcription initiation factor IIA subunit 1-like isoform X1 | Hill | OrgC |
| LOC124342780 | transforming growth factor-beta-induced protein ig-h3-like | Hill | OrgC |
| LOC124337916 | transitional endoplasmic reticulum ATPase-like isoform X1 | Hill | OrgC |
| LOC124336851 | tudor domain-containing 6-like | Hill | OrgC |
| LOC124310854 | tyrosine-protein phosphatase 99A-like | Hill | OrgC |
| LOC124341834 | UDP-glucosyltransferase 2-like | Hill | OrgC |
| LOC124341836 | UDP-glucosyltransferase 2-like | Hill | OrgC |
| LOC124344435 | UDP-glucosyltransferase 2-like | Hill | OrgC |
| LOC124341270 | UNC93-like protein | Hill | OrgC |
| LOC124314862 | uncharacterized protein KIAA1143 homolog | Hill | OrgC |
| LOC124314721 | uncharacterized protein LOC124314721 isoform X1 | Hill | OrgC |
| LOC124316145 | uncharacterized protein LOC124316145 isoform X1 | Hill | OrgC |
| LOC124327540 | uncharacterized protein LOC124327540 isoform X1 | Hill | OrgC |
| LOC124342070 | uncharacterized protein LOC124342070 | Hill | OrgC |
| LOC124342144 | uncharacterized protein LOC124342144 | Hill | OrgC |
| LOC124342628 | uncharacterized protein LOC124342628 | Hill | OrgC |
| LOC124342817 | uncharacterized protein LOC124342817 | Hill | OrgC |
| LOC124342892 | uncharacterized protein LOC124342892 | Hill | OrgC |
| LOC124343590 | uncharacterized protein LOC124343590 | Hill | OrgC |
| LOC124350663 | uncharacterized protein LOC124350663 | Hill | OrgC |
| LOC124350679 | uncharacterized protein LOC124350679 isoform X1 | Hill | OrgC |
| LOC124342806 | uncharacterized serine-rich protein C215.13-like | Hill | OrgC |
| LOC124316173 | vascular endothelial growth factor receptor 1-like | Hill | OrgC |
| LOC124316498 | vascular endothelial growth factor receptor 3-like isoform X1 | Hill | OrgC |
| LOC124316458 | WSC domain-containing protein 2-like | Hill | OrgC |
| LOC124316763 | WSC domain-containing protein 2-like | Hill | OrgC |
| LOC124327665 | zinc finger MYM-type protein 1-like | Hill | OrgC |
| LOC124342568 | zinc finger protein 229-like isoform X3 | Hill | OrgC |
| LOC124316314 | zinc finger protein 660-like isoform X1 | Hill | OrgC |
| LOC124342568 | zinc finger protein rotund-like isoform X1 | Hill | OrgC |
| LOC124335734 | zinc metalloproteinase nas-13-like | Hill | OrgC |
| LOC124349671 | endocuticle structural glycoprotein SgAbd-3-like isoform X1 | SC | OrgC |
| LOC124348872 | hatching enzyme 1.2-like | SC | OrgC |
| LOC124310908 | synaptogenesis protein syg-2-like isoform X1 | SC | OrgC |
| LOC124348859 | T-box transcription factor TBX2-like isoform X1 | SC | OrgC |
| LOC124348859 | T-box transcription factor TBX2b-like isoform X3 | SC | OrgC |
| LOC124326489 | uncharacterized protein LOC124326489 | SC | OrgC |
| LOC124344527 | uncharacterized protein LOC124344527 | SC | OrgC |
| LOC124349363 | uncharacterized protein LOC124349363 isoform X1 | SC | OrgC |
| LOC124316511 | alpha-ketoglutarate-dependent dioxygenase alkB homolog 7%2C mitochondrial-like | Hill | P |
| LOC124315722 | aminoacylase-1-like | Hill | P |
| LOC124314224 | collagen alpha-1(XVIII) chain-like isoform X1 | Hill | P |
| LOC124315050 | lysine-specific demethylase 4C-like isoform X1 | Hill | P |
| LOC124315040 | neuropeptide F receptor-like | Hill | P |
| LOC124313863 | protein abrupt-like | Hill | P |
| LOC124318419 | U6 snRNA-associated Sm-like protein LSm8 | Hill | P |
| LOC124314010 | uncharacterized protein LOC124314010 isoform X1 | Hill | P |
| LOC124320214 | uncharacterized protein LOC124320214 | Hill | P |
| LOC124313864 | voltage-dependent L-type calcium channel subunit beta-2-like isoform X1 | Hill | P |
| LOC124315571 | zinc finger protein 277-like | Hill | P |

**Table S2: South Center Fst Outlier GO term analysis output from PantherDB webtool.**

##Analysis Type:	PANTHER Overrepresentation Test (Released 20230705)

##Annotation Version and Release Date:	PANTHER version 17.0 Released 2022-02-22

##Analyzed List:	SC_outliers_fst_panther.tsv

##Reference List:	Daphnia pulex (all genes in database)

##Test Type:	FISHER

##Correction:	FDR

PANTHER GO-Slim Molecular Function	Daphnia pulex - REFLIST (30047)	SC_outliers_fst_panther.tsv (120)	SC_outliers_fst_panther.tsv (expected)	SC_outliers_fst_panther.tsv (over/under)	SC_outliers_fst_panther.tsv (fold Enrichment)	SC_outliers_fst_panther.tsv (raw P-value)	SC_outliers_fst_panther.tsv (FDR)

monocarboxylic acid transmembrane transporter activity (GO:0008028)	13	2	.05	+	38.52	1.59E-03	3.00E-02

phosphatidylinositol bisphosphate binding (GO:1902936)	22	3	.09	+	34.14	1.32E-04	3.55E-03

phosphatidylinositol phosphate binding (GO:1901981)	28	3	.11	+	26.83	2.54E-04	6.17E-03

transcription coactivator activity (GO:0003713)	33	3	.13	+	22.76	3.98E-04	9.21E-03

phosphatidylinositol binding (GO:0035091)	39	3	.16	+	19.26	6.29E-04	1.39E-02

neurotransmitter receptor activity (GO:0030594)	57	3	.23	+	13.18	1.78E-03	3.24E-02

neurotransmitter binding (GO:0042165)	57	3	.23	+	13.18	1.78E-03	3.13E-02

phospholipid binding (GO:0005543)	66	3	.26	+	11.38	2.66E-03	4.51E-02

G protein-coupled receptor activity (GO:0004930)	94	4	.38	+	10.65	6.45E-04	1.37E-02

signaling receptor activity (GO:0038023)	279	10	1.11	+	8.97	2.56E-07	3.26E-05

molecular transducer activity (GO:0060089)	280	10	1.12	+	8.94	2.64E-07	2.24E-05

transmembrane signaling receptor activity (GO:0004888)	168	6	.67	+	8.94	7.16E-05	2.28E-03

transcription coregulator activity (GO:0003712)	113	4	.45	+	8.86	1.25E-03	2.54E-02

ion channel activity (GO:0005216)	145	4	.58	+	6.91	3.01E-03	4.79E-02

transcription regulator activity (GO:0140110)	438	12	1.75	+	6.86	2.59E-07	2.63E-05

RNA polymerase II transcription regulatory region sequence-specific DNA binding (GO:0000977)	330	9	1.32	+	6.83	9.00E-06	5.73E-04

DNA-binding transcription factor activity, RNA polymerase II-specific (GO:0000981)	304	8	1.21	+	6.59	3.69E-05	1.34E-03

transcription cis-regulatory region binding (GO:0000976)	350	9	1.40	+	6.44	1.42E-05	8.03E-04

transcription regulatory region nucleic acid binding (GO:0001067)	350	9	1.40	+	6.44	1.42E-05	7.23E-04

sequence-specific DNA binding (GO:0043565)	391	10	1.56	+	6.40	4.88E-06	3.55E-04

sequence-specific double-stranded DNA binding (GO:1990837)	361	9	1.44	+	6.24	1.80E-05	8.34E-04

DNA-binding transcription factor activity (GO:0003700)	324	8	1.29	+	6.18	5.71E-05	1.94E-03

double-stranded DNA binding (GO:0003690)	393	9	1.57	+	5.73	3.45E-05	1.35E-03

ion transmembrane transporter activity (GO:0015075)	242	5	.97	+	5.17	3.14E-03	4.84E-02

DNA binding (GO:0003677)	579	11	2.31	+	4.76	2.48E-05	1.05E-03

transmembrane transporter activity (GO:0022857)	407	7	1.63	+	4.31	1.37E-03	2.69E-02

transporter activity (GO:0005215)	460	7	1.84	+	3.81	2.70E-03	4.44E-02

nucleic acid binding (GO:0003676)	1035	14	4.13	+	3.39	7.56E-05	2.14E-03

organic cyclic compound binding (GO:0097159)	1310	16	5.23	+	3.06	7.43E-05	2.22E-03

heterocyclic compound binding (GO:1901363)	1291	15	5.16	+	2.91	2.16E-04	5.51E-03

binding (GO:0005488)	2874	31	11.48	+	2.70	2.46E-07	4.17E-05

molecular_function (GO:0003674)	5854	51	23.38	+	2.18	9.53E-09	4.85E-06

Unclassified (UNCLASSIFIED)	24193	69	96.62	-	.71	9.53E-09	2.43E-06

**Table S3: Hill Lake Organic Carbon GO output**

##Analysis Type:	PANTHER Overrepresentation Test (Released 20230705)

##Annotation Version and Release Date:	PANTHER version 17.0 Released 2022-02-22

##Analyzed List:	Hill_OrgC_outliers_panther.tsv

##Reference List:	Daphnia pulex (all genes in database)

##Test Type:	FISHER

##Correction:	FDR

PANTHER GO-Slim Molecular Function	Daphnia pulex - REFLIST (30047)	Hill_OrgC_outliers_panther.tsv (58)	Hill_OrgC_outliers_panther.tsv (expected)	Hill_OrgC_outliers_panther.tsv (over/under)	Hill_OrgC_outliers_panther.tsv (fold Enrichment)	Hill_OrgC_outliers_panther.tsv (raw P-value)	Hill_OrgC_outliers_panther.tsv (FDR)

sequence-specific double-stranded DNA binding (GO:1990837)	361	6	.70	+	8.61	7.47E-05	1.27E-02

DNA-binding transcription factor activity (GO:0003700)	324	5	.63	+	7.99	4.31E-04	3.66E-02

sequence-specific DNA binding (GO:0043565)	391	6	.75	+	7.95	1.15E-04	1.46E-02

double-stranded DNA binding (GO:0003690)	393	6	.76	+	7.91	1.18E-04	1.20E-02

RNA polymerase II transcription regulatory region sequence-specific DNA binding (GO:0000977)	330	5	.64	+	7.85	4.68E-04	3.40E-02

transcription cis-regulatory region binding (GO:0000976)	350	5	.68	+	7.40	6.09E-04	3.87E-02

transcription regulatory region nucleic acid binding (GO:0001067)	350	5	.68	+	7.40	6.09E-04	3.44E-02

DNA binding (GO:0003677)	579	6	1.12	+	5.37	9.02E-04	4.17E-02

catalytic activity (GO:0003824)	3091	15	5.97	+	2.51	6.19E-04	3.15E-02

molecular_function (GO:0003674)	5854	28	11.30	+	2.48	7.69E-07	1.96E-04

Unclassified (UNCLASSIFIED)	24193	30	46.70	-	.64	7.69E-07	3.92E-04
